# Notch Controls Multiple Pancreatic Cell Fate Regulators Through Direct Hes1-mediated Repression

**DOI:** 10.1101/336305

**Authors:** Kristian H. de Lichtenberg, Philip A. Seymour, Mette C. Jørgensen, Yung-Hae Kim, Anne Grapin-Botton, Mark A. Magnuson, Nikolina Nakic, Jorge Ferrer, Palle Serup

## Abstract

Notch signaling and its effector Hes1 regulate multiple cell fate choices in the developing pancreas, but few direct target genes are known. Here we use transcriptome analyses combined with chromatin immunoprecipitation with next-generation sequencing (ChIP-seq) to identify direct target genes of Hes1. ChIP-seq analysis of endogenous Hes1 in 266-6 cells, a model of multipotent pancreatic progenitor cells, revealed high-confidence peaks associated with 354 genes. Among these were genes important for tip/trunk segregation such as *Ptf1a* and *Nkx6-1*, genes involved in endocrine differentiation such as *Insm1* and *Dll4*, and genes encoding non-pancreatic basic-Helic-Loop-Helix (bHLH) factors such as *Neurog2* and *Ascl1*. Surprisingly, we find that Hes1 binds a large number of loci previously reported to bind Ptf1a, including a site downstream of the *Nkx6-1* gene. Notably, we find a number of Hes1 bound genes that are upregulated by γ-secretase inhibition in pancreas explants independently of *Neurog3* function, including the tip progenitor/acinar genes; *Ptf1a, Gata4, Bhlha15*, and *Gfi1*. Together, our data suggest that Notch signaling suppress the tip cell fate by Hes1-mediated repression of the tip-specific gene regulatory network module that includes transcriptional regulators such as Ptf1a, Gata4, Mist1, and Gfi1. Our data also uncover new molecular targets of Notch signaling that may be important for controlling cell fate choices in pancreas development.

## Introduction

Notch signaling plays a pivotal role in pancreas development by regulating progenitor proliferation and cell fate decisions and thus ensures the precise and timely regulation of the cell number of each lineage, which is ultimately crucial for proper organ function [1]. The juxtacrine Notch signaling pathway is characterized by signal sending from one cell via the DSL (Delta/Serrate/LAG-2)-family of ligands. In mammals four ligands (Dll1, Dll4, Jag1, and Jag2) are capable of transactivating Notch receptors on neighboring cells Transactivation involves ligand ubiquination catalyzed by Mib1, which is required for ligand endocytosis. Ligand endocytosis is thought to exert a pulling force on Notch receptors that allow cleavage by ADAM10 and the γ-secretase complex [2-5]. This leads to release of the Notch Intracellular domain (NICD), translocation to the nucleus for activation of transcription through the obligate DNA binding partner Rbpj and the co-activator Maml1. Prominent among pancreatic Notch target genes is *Hes1*, which encodes a transcriptional repressor of the Hairy and Enhancer-of-split (Hes)-family of bHLH proteins [6].

The pancreatic buds evaginate from the primitive gut tube around E9.0 in mouse development and is comprised of multipotent progenitor cells (MPCs) that give rise to the entire pancreas epithelium. Dll1-induced Notch activity in the MPCs acts to limit early endocrine differentiation through Hes1-mediated repression of the proendocrine gene *Neurog3* and to maintain normal MPC proliferation [6-9]. Hes1 and Neurog3 proteins are typically detected in a mutually exclusive pattern and this is likely a reflection of the lateral inhibition feedback mechanism. Lateral inhibition posits that a trunk progenitor undergoing endocrine differentiation will express more Dll1 than its neighbors, leading to Notch activation, upregulation of Hes1, and repression of Neurog3 (and Dll1) in the neighboring cells, which consequently will remain in a progenitor state. However, while some aspects of Notch-mediated lateral inhibition are likely to operate in the pancreas the complete mechanism is likely to be more complicated [7, 10]. Indeed, recent work suggests that EGF signaling is involved in initiating the downregulation of Hes1 that leads to derepression of *Neurog3* and endocrine differentiation [11].

Beginning around E10, MPCs progressively segregate into acinar cell fated tip progenitors and duct/endocrine fated trunk progenitors that can be followed by the segregation of a set of molecular markers. Initially, MPCs are marked by co-expression of several transcription factors whose expression later diverge to either the tip lineage (Ptf1a and Gata4) or the trunk lineage (Nkx6-1, Sox9, Hnf1β, Onecut1, and Gata6; reviewed in [12]). These transcription factors engage in a complex gene regulatory network (GRN) whose wiring is likely to be dynamic and change over time. For example, Ptf1a is necessary for proper Nkx6-1 expression in MPCs whereas later it will repress Nkx6-1 to establish a Ptf1a^+^Gata4^+^ tip/acinar fate over a Nkx6-1^+^Hnf1^+^Sox9^+^ trunk/bipotent progenitor fate. Conversely, forced expression of Nkx6-1 in MPCs represses Ptf1a expression and tip fate [13, 14]. In addition to its early function as a suppressor of endocrine differentiation, Notch signaling is also important for proper allocation of tip and trunk fates. Forced expression of N1ICD prevents the cells from adopting a tip fate [13, 15, 16] and N1ICD has been suggested to act by directly activating Nkx6-1 expression [17] and to suppress Ptf1a expression and/or function indirectly via induction of Hes1 [18, 19]. Conversely, conditional *Mib1* mutants that have compromised Notch ligand activity show a complete conversion of trunk cells to a tip fate and to a lesser degree the same phenotype is seen in conditional Hes1 mutants and after expression of dominant negative Maml1 [17, 20]. Thus, forced expression of Ptf1a phenocopies Notch loss-of-function mutations and forced expression of Nkx6-1 phenocopies Notch gain-of-function mutations but exactly how Notch interfaces with the above mentioned GRN, and at how many nodes, remains unclear.

In the present study, we have identified direct target genes of Hes1 and examined gene expression changes and pancreatic cell fate choices regulated by Notch signaling. We found 354 Hes1 bound genes in 266-6 cells, a model of MPCs previously used to define Ptf1a target genes [14]. Most notable among these were genes important for tip/trunk segregation such as *Ptf1a* and *Nkx6-1* as well as genes involved in regulating endocrine differentiation such as *Insm1* and *Dll4*. Surprisingly, we find Hes1 to bind a large number of distal regulatory elements previously reported to bind Ptf1a, including a site in the *Nkx6-1* gene. Notably, we find multiple Hes1 bound genes, including tip progenitor/acinar markers; *Ptf1a, Gfi1*, and *Gata4*, that are deregulated by treatment with DAPT, a γ-secretase inhibitor (GSI). Lastly, we find the gene *Cbfa2t3*, encoding a transcriptional corepressor, to be bound by Hes1 and upregulated in the explant models after loss of Notch/Hes1 function. Together, our data suggest that Notch signaling regulate tip/trunk patterning by Hes1-mediated repression of multiple tip/acinar-specific transcriptional regulators and uncover new molecular targets of Notch signaling that may be important for controlling cell fate choices in pancreas development.

## RESULTS

### Transcriptomes of Hes1^+^ and Hes1^−^ pancreatic progenitor populations

In order to generate a comprehensive map of differentially expressed genes in embryonic mouse pancreas cell types differing in their Hes1 expression status, we first performed microarray-based gene expression profiling of purified E15.5 pancreas cell populations. As outlined in Figure 1A, we used a combination of Dolichos Biflorus Agglutinin (DBA) labeling and a double transgenic *Neurog3*-RFP; *Hes1*-GFP mouse line to isolate Hes1^+^ bipotent trunk progenitors (GFP^+^RFP^−^DBA^+^) as well as presumptive early (GFP^+^ RFP^+^DBA^+^) and late (GFP^+^RFP^+^DBA^−^) Hes1^−^ endocrine precursor cells from dissociated E15.5 pancreata via fluorescence activated cell sorting (FACS). In parallel FACS experiments we isolated Hes1^−^ tip/acinar precursors cells (YFP^+^) from dissociated E15.5 *Ptf1a*^Yfp/+^ pancreata (Figure 1B). We then subjected all four cell populations to gene expression analysis by microarray analysis. Principal Component Analysis using hierarchical clustering of the individual samples revealed close grouping of the different biological replicates indicating high reproducibility between experiments (Figure 1C). The heatmap shown in Figure 1D reveals that the presumptive early and late endocrine precursor populations (Hes1-GFP^+^Neurog3-RFP^+^DBA^+^ and Hes1-GFP^+^Neurog3-RFP^+^DBA^−^) share gene expression profiles and we found no significant differentially expressed genes between the two populations (log2 fold change >1.5 and Benjamini-Hochberg (BH) adjusted p-value <0.1), thus, for subsequent analyses we pooled the two endocrine precursor populations (GFP^+^RPF^+^DBA^+/−^). Differential expression analysis comparing the endocrine precursors to the GFP^+^RPF^−^DBA^+^ trunk progenitor population revealed a cluster of genes (Cluster 5) with increased expression and that included many known endocrine lineage markers such as *Neurog3, Neurod1, Insm1, Glucagon* and *Chga* (Figure 1D, Sup. Figure 1A, and Supplemental Table 1). Gene Set Enrichment Analysis (GSEA) of the pooled endocrine precursor populations (GFP^+^RFP^+^DBA^+/−^) compared to both the YFP^+^ acinar and GFP^+^RFP^−^DBA^+^ trunk progenitors, using the MSigDb:C2 database revealed strong enrichment of GLIS3 target genes (KANG GLIS3 TARGET) and the Reactome gene sets “Regulation of Insulin Secretion” and “Integration of Energy Metabolisms” (Supplemental Figure 1B), together signifying that the GFP^+^RFP^+^DBA^+/−^ populations represent endocrine precursor cells, and likely contain some mature endocrine cells as well. Consistent with previous work [21], differential expression analysis of the YFP^+^ acinar population compared to the GFP^+^RPF^−^DBA^+^ bipotent/trunk progenitors revealed that acinar cells share expression of more genes with bipotent progenitors than do endocrine progenitors Figure 1D), but also display significantly upregulated expression of classic acinar markers including *Rbpjl* (11.0-fold), *Cela1* (24.9-fold), *Ptf1a* (5.8-fold), *Gata4* (3.4-fold), *Gfi1* (7.4-fold) and *Cpa2* (12.2-fold) as shown on the volcano plot in Supplemental Figure 2 [22]. GSEA found the acinar progenitors to be enriched for Myc target genes (Schumacher Myc Targets Up), which is reassuring since Myc is a known acinar marker [23] and accordingly its transcript is enriched in the YFP^+^ acinar population. A second gene set, “Sotiriou Breast Cancer Grade 1 vs 3 Up”, is mainly containing genes associated with cell cycle and proliferation in line with the marked proliferation seen in tip/acinar cells at this stage of development [24]. Also, we found “Zhang TLX targets 36hr Down” to be enriched, a gene-set containing genes down-regulated in neural stem cells after *Nr2e1* knock-out (previously knowns as TLX), also including cell cycle and proliferation markers such as *Mki67*.

**Figure 1.**
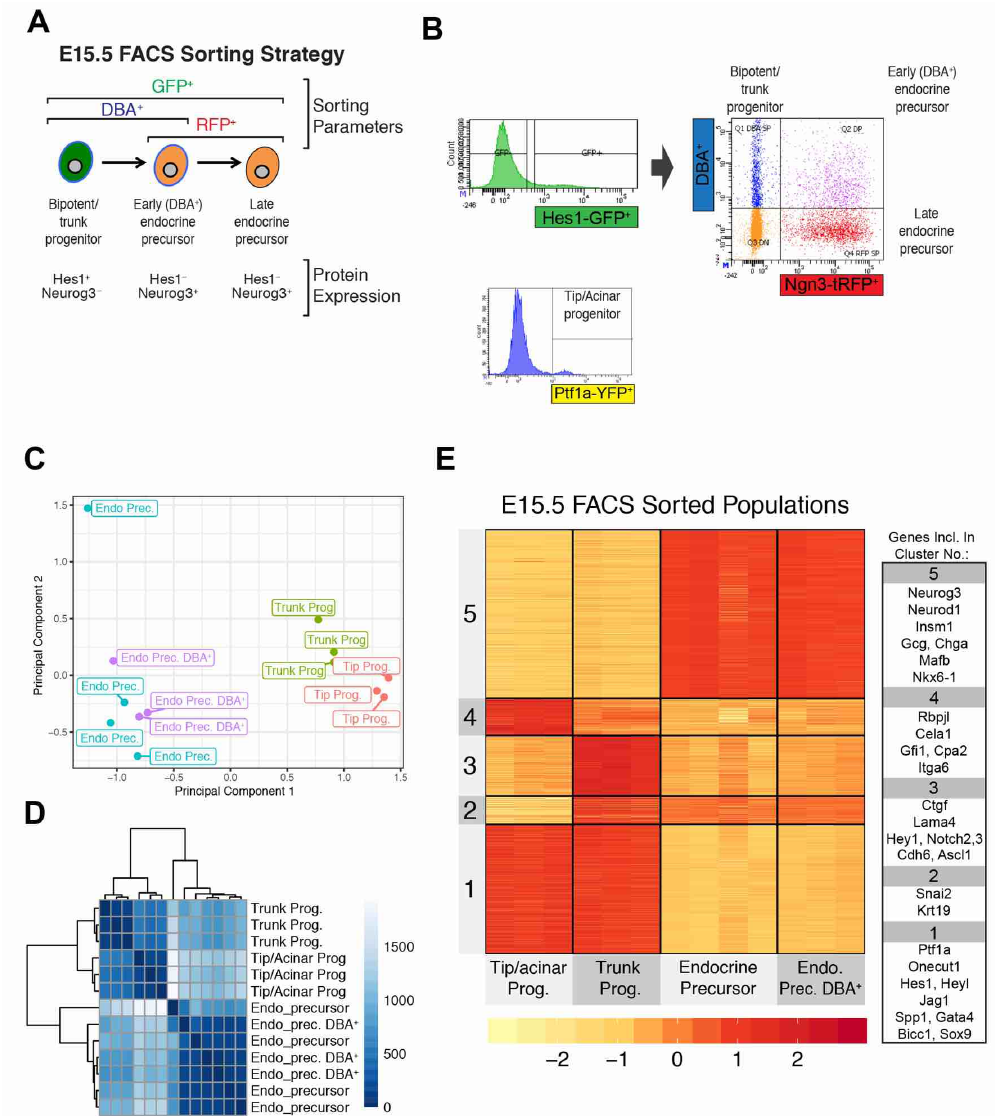
FACS sorting and microarray gene expression analysis of E15.5 pancreas populations. A) Schematic of experimental setup with expected reporter status and protein expression. B) FACS setup of Hes1-GFP;Neurog3-RFP mice stained with DBA lectin, on the left a representative FACS histogram of Hes1-GFP^+^ cells, which were then gated for Hes1-GFP^+^DBA^+^ (bipotent/ trunk progenitors), Hes1-GFP^+^DBA^+^Neurog3-RFP^+^ (endocrine precursors-DBA^+^) and Hes1-GFP^+^DBA-Neurog3-RFP^+^ (endocrine precursors DBA^−^). On the right representative histogram from FACS analysis of Ptf1a-YFP^+^ cells, sorted to obtain the acinar population. C) MultiDimensional Scaling analysis of E15.5 FACS sorted microarray samples. D) Hierarchical clustering analysis of E15.5 FACS sorted microarray samples. E) Heatmap of differentially expressed genes in any comparison clustered by kmeans, scaled by row. On the right representative examples of genes from each cluster.

**Figure 2.**
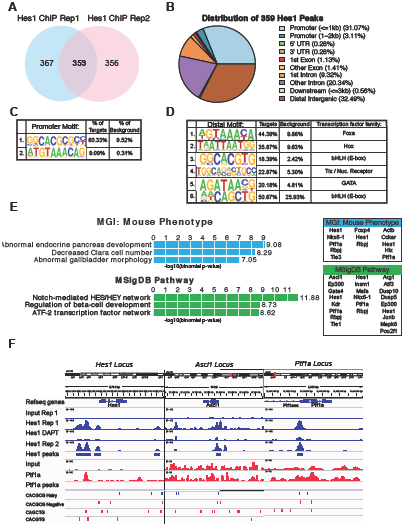
Chip-seq of Hes1 in 266-6 cell line. A) Venn diagram showing the overlap in bold between the two ChIP-seq replicates. B) Distribution of Hes1 peaks by MACS2 peak finding on genomic features. C) Top 2 hits from Homer de novo motif finding at the promoter region (2kp upstream and 0.5kb downstream a transcription start site), the Hes1 motif and at lower frequency a Foxa family motif. D) Top enriched Homer de novo motif finding analysis on Hes1 binding sites distal to promoters (more than 2kb upstream or 0.5kb downstream transcription start site) with associated transcription factor families as indicated. E) GREAT enrichment of annotation analyses of the Hes1 peaks: MGI Mouse Phenotypes and MSigDB Pathway, and in the box on the right are the genes within the respective terms. F) IGV Screenshot of Hes1 ChIP-seq enrichment at indicated loci, with Hes1 ChIP-seq replicate 1 and 2 trace in blue, controls Input and Hes1 ChIP-seq in DAPT (Notch inhibited conditions) in black. RefSeq gene track and below that Hes1 peaks found by MACS2 and motifs as indicated on the left.

Conversely, and as expected, we find trunk markers upregulated in the GFP^+^RFP^−^DBA^+^ population: Sox9 (2.4-fold), Hnf1b (2.2-fold), Krt19 (3.2-fold), Nkx6-1 (2.9-fold), Spp1 (3.2-fold), Bicc1 (3.8-fold) and GSEA analyses yielded signatures largely comprised of extracellular matrix related genes including several collagens, laminins, and cadherins shown in Sup. Figure 2. GSEA of the signatures enriched in the bipotent/trunk population compared to the endocrine precursors largely contained cell cycle and proliferation markers in gene sets such as “Gobert Oligodendrocyte Differentiation Up” and “Zhang Cell Cycle” (Sup. Figure 1C) highlighting the proliferative differences between trunk cells and the post-mitotic endocrine precursors [25]. Together this data set provides a comprehensive gene expression atlas of distinct progenitor populations with different Hes1 expression profiles. In the following sections, we map additional data sets from Hes1 ChIP-seq experiments and microarray analyses of GSI treated explants onto this gene expression atlas, in order to define Hes1 target genes.

### Hes1 Chromatin Immunoprecipitation in 266-6 cell line reveals potential direct target genes

To investigate the genome-wide binding of Hes1 to find direct target genes in the mouse pancreas, we performed chromatin immunoprecipitation with next generation sequencing (ChIP-seq) of Hes1 in the acinar carcinoma cell line 266-6. This cell line, derived from an Ela-SV40 large T transgenic mouse [26] is the best available model of mouse multipotent pancreas progenitor cells available with co-expression of Pdx1, Onecut1, Ptf1a, and Nkx6-1 [14]. We observed that Notch signaling is active in these cells and that they respond to γ-secretase inhibition by reduced N1-ICD and Hes1 protein levels (Figure 4). We performed ChIP-seq of Hes1 and did peak calling using MACS2 and among two biological replicates we found 359 common peaks of which 1/3 were found to be in promoter regions within 2kb of the Transcription Start Site (TSS), 1/3 located in introns and exons and the remaining 1/3 in distal intergenic regions (Figure 2A and B). De novo motif search on Hes1 peaks close to promoters using Homer found an extended motif GGCACGCGCC motif (Figure2C), encompassing the classical C-site (underlined) described for Hairy proteins [27]. The second most enriched motif is of the Foxa family although at a much lower frequency. De novo motif search at the Hes1-bound loci distal to promoters revealed a Foxa motif (Figure 2D), a Hox-related site similar to the Pdx1 binding site, an extended bHLH motif GGCACGTG containing the classical Hairy B-site motif (underlined, [27]), Nuclear receptor (Tlx), a Gata motif, and another bHLH motif CAGCTG, corresponding to the most abundant Ptf1a-bound site in the developing spinal cord, which is also the preferred recognition sequence for Ascl1 and Atoh1 proteins [28-31].

GREAT annotation and gene ontology (GO) analysis of the peaks revealed enrichment of the MGI Mouse Phenotype “Abnormal endocrine pancreas development” driven by the genes *Hes1, Nkx6-1, Ptf1a, Rbpj and Tle3* as indicated in the right panel of Figure2E. “Decreased Clara cell number” were only given by three genes, *Foxp4, Hes1* and *Rbpj,* which are all expressed and relevant in the pancreas. We also found significant enrichment in the MSigDB pathway database through GREAT, with “Notch-mediated HES/HEY network” to be most enriched containing following genes bound: *Ascl1, Ep300, Gata4, Hes1, Kdr, Ptf1a, Rbpj and Tle1*.

Tracks of the Hes1 ChIP-seq tracks reveal three distinct Hes1 peaks upstream of the *Hes1* gene itself confirming Hes1 binding to its own promoter in a pancreatic cell compared to negative controls Input and Hes1 ChIP-seq on DAPT treated cells which inhibits Notch signaling and thereby Hes1 expression (Figure2 and additional tracks in Sup. Figure 3). Interestingly, *Neurog2* which is not expressed in the pancreas but is a Hes1 target in neuronal stem cells and in mouse embryonic stem cells [32] was found bound, whereas we did not find the most prominent expected Hes1 target in the pancreas, *Neurog3* [6, 33, 34], emphasizing the limitations of the acinar cell-derived 266-6 cell line (see discussion).

**Figure 3.**
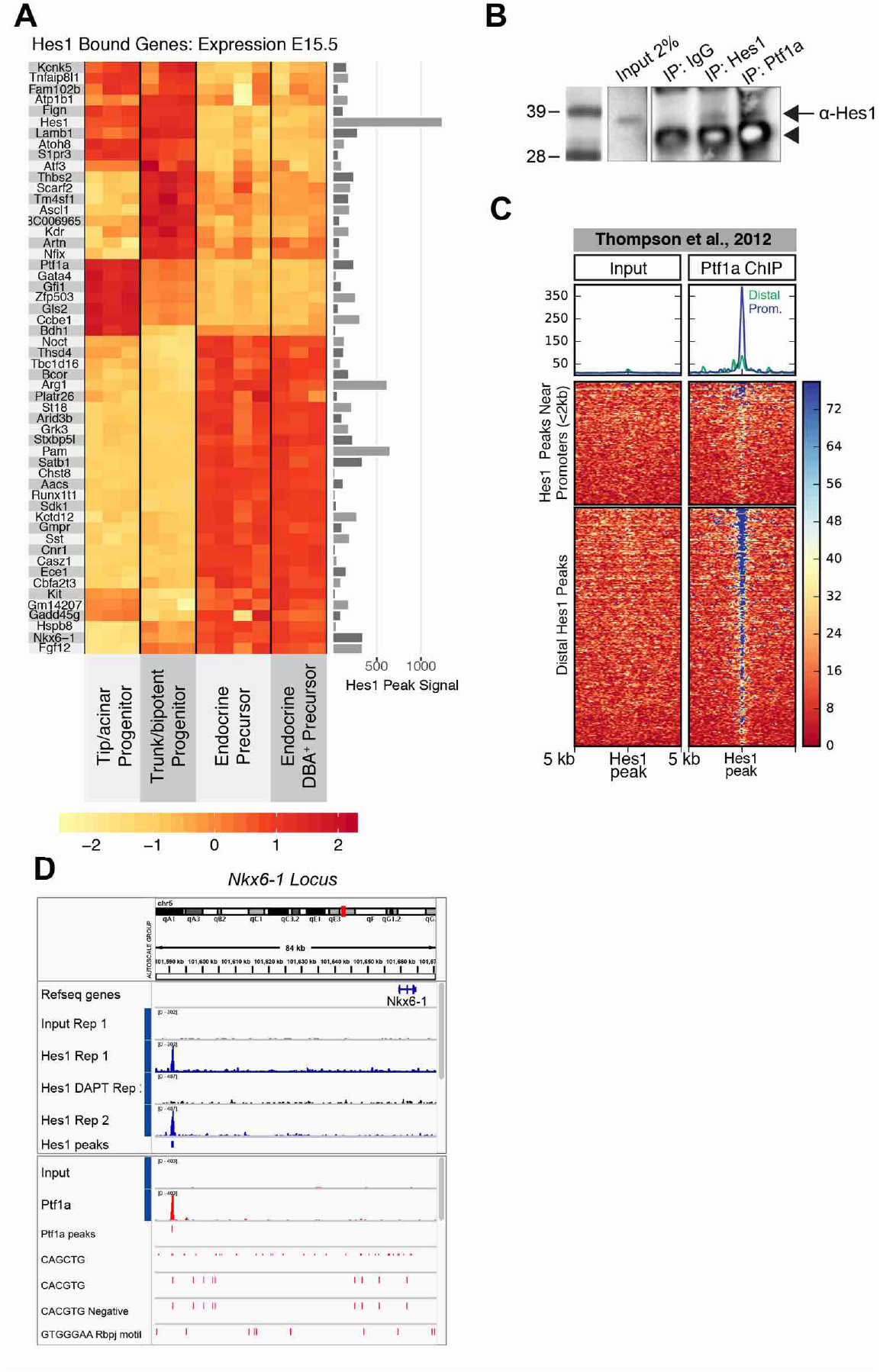
Hes1 ChIP-seq overlap with Ptf1a-ChIP and overlap with E15.5 FACS sorted pancreas population expression data. A) Heatmap of expression profiles from E15.5 pancreas populations on Hes1 bound genes. Enrichment under the Hes1 peak shown with bars on the right B) Western blot of Immunoprecipiation with IgG control, Hes1 positive control and Ptf1a antibodies, blotting for Hes1 protein. Arrow indicating Hes1 band. Arrow head indicating IgG light chain from immunoprecipitation procedure. C) Heatmap of Ptf1a binding to Hes1 peaks +/−5kb, divided by proximity to promoter (<2kb) on upper heatmap shown with green trace in summary line plot. Lower heatmap with distal Hes1 bound sites summarized as blue trace in line plot. D) ChIP-seq tracks at the Nkx6-1 locus, with Hes1 ChIP-seq tracks in blue, controls in black and Ptf1a ChIP-seq in red (from Thompson et al., 2012). Hes1 and Ptf1a peaks as indicated and coloured in correspondence to tracks and below motifs for Hes1, Ptf1a and Rbpj as indicated on the left.

### Ptf1a binds to the same genomic regions as Hes1 and forms a stable complex with Hes1

To evaluate how Hes1 is participating the cell fate choices in pancreas development we analyzed *in vivo* gene expression profiles of the Hes1 bound genes, in our E15.5 FACS sorted pancreas populations. We visualized the expression levels of Hes1 bound genes that are differentially expressed between endocrine, acinar and trunk progenitors in a heatmap (Figure 3A). If genes that are bound by the transcriptional repressor Hes1 in 266-6 cells show highest expression in trunk progenitors, where Hes1 is also present, they must have additional activating inputs, e.g. via N-ICD or a lineage-specific transcription factor, or be oscillating as is the case for *Ascl1* and *Hes1* in neural progenitors [35]. The genes that are highly expressed in the endocrine progenitors, and bound by Hes1 in 266-6 cells, are likely to be derepressed as Hes1 expression is lost upon commitment to the endocrine lineage. Such genes include *Somatostatin (Sst), Insm1* and the previously described direct Hes1 target, *Gadd45g*, involved in growth arrest [32].

Serendipitously, we noted binding of Hes1 to one of two distal regulatory elements surrounding *Nkx6-1* previously shown by Thompson and colleagues to be bound by Ptf1a (Figure 3D), which prompted us to assess Ptf1a enrichment on our Hes1 peaks from published Ptf1a ChIP-seq data also made in 266-6 cell line [14]. We found extensive Ptf1a binding on distal Hes1 peaks but rarely on promoters bound by Hes1 (Figure 3C), consistent with the previously reported preferential binding of Ptf1a to distal sites [36] and the above mentioned *de novo* motif finding analysis that identified the Ptf1a site CATCTG. In addition to *Nkx6-1*, examples of Hes1 and Ptf1a binding to the same locus includes the *Hes1* locus where Ptf1a binds within one of three major Hes1 peaks, the *Acvrl1, Dll4/GM14207*, and *Gfi1* loci, the latter which is bound in the promoter region (Sup. Figure 3). The closeness of the Hes1 and Ptf1a binding sites suggest that Hes1 and Ptf1a may interact directly. Interaction between overexpressed, epitope-tagged Hes1 and Ptf1a has previously been reported by Ghosh and Leach (Ghosh and Leach, 2006) and we can now show interaction between endogenous levels of these two proteins in 266-6 cells by co-immunoprecipitation (Figure 3B). Together with our ChIP-seq data this suggests a novel mechanism of transcriptional co-regulation by Hes1 and Ptf1a at distal regulatory sites. Similarly, We also found extensive co-binding between Hes1 and Foxa2, consistent with the reported co-binding of Ptf1a and Foxa2 and our identification of Foxa sites by *de novo* motif search. However, Foxa2 co-binding included more than Ptf1a co-bound sites and also included sites co-bound by Rbpj, but not Ptf1a and Rbpjl, suggesting that these may represent Notch response elements (Sup. Figure 7).

### Notch Inhibition Increases Expression of Acinar Fate Determinants

Since loss of Hes1 and Notch signalling drives progenitors to the endocrine lineage, we set up a pancreas explant system with crosses of heterozygous *Neurog3*^*tTA/+*^, a knock-in allele which makes the homozygote *Neurog3*^*tTA/tTA*^ embryos deficient in *Neurog3* (hereafter *Neurog3*-null). Hereby endocrine differentiation is impeded and we can study the Neurog3-independent role of Notch signalling by DAPT inhibition of γ-secretase. E12.5 wildtype and *Neurog3*-null pancreata were explanted on fibronectin, grown for 3 days, and then treated with vehicle control (0.1% DMSO) or DAPT for 24h. RNA was isolated and subjected to Agilent microarray analysis. With exception of the wildtype vehicle control, the samples clustered by similarity according to genotype and treatment In a MultiDimensional scaling (MDS) plot (Sup. Figure 5). This was also reflected in the differential expression analysis by Limma showing only three RefSeq genes differentially expressed between wildtype+DAPT versus wildtype+DMSO (BH adj.p-value threshold <0.1 and log2 fold change >1), namely *Cck, Espn* and *Neurog3* (Sup. Figure 4). The robust upregulation of Neurog3 despite the variability in the wildtype DMSO samples confirms that Notch signalling is suppressed after 24h DAPT treatment. We found 104 differentially expressed RefSeq-genes when comparing DAPT treated *Neurog3*-null and wildtype samples. As expected, endocrine genes (defined as highly expressed in the sorted E15.5 GFP^+^RFP^+^DBA^+/−^ endocrine precursor populations) showed higher expression in the wildtype than Neurog3-null (Sup. Figure 5). All the islet hormone genes and *Neurog3* were included in this cluster (Sup. Figure 4). When comparing DAPT and vehicle treated Neurog3-null explants, we found 233 upregulated and 65 downregulated genes, the latter including: *Reg3b, Fgf6, Fgf20* and trunk/duct marker *Spp1* (Osteopontin) (Figure 4B and Sup. Figure 4). We reasoned that the downregulated genes could be direct Notch target genes, but did not find Notch related signatures or Rbpj-motifs enriched by GSEA (data not shown), however more in-depth analysis could reveal a direct link to Notch target genes.

**Figure 4.**
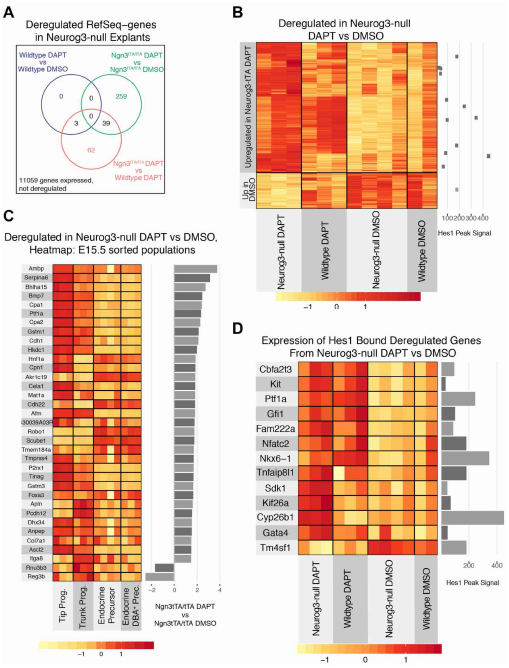
Deregulated genes in DAPT-treated Neurog3-null pancreas explants at E12.5+4d. A) Venn diagram of differentially expressed genes in comparisons as indicated. Neurog3-null (Neurog3^tTA/tTA^) DMSO versus wildtype DMSO comparison yielded no differentially expressed genes and was not included. B) Scaled expression of 298 differentially expressed genes between Neurog3-null DAPT and Neurog3-null DMSO treated with corresponding Hes1 peaks for those genes that are bound and the enrichment under the peak in the scatterplot on the right. C) Bar plot of the top genes by log2 fold change between Neurog3-null DAPT and Neurog3-null DMSO, with heatmap showing the expression of corresponding gene in E15.5 FACS sorted pancreas populations. D) Hes1 bound by ChIP-seq in 266 which are also deregulated genes in Neurog3-null DAPT and Neurog3-null DMSO. Hes1 peak enrichment as bar graph and scaled expression in heatmap.

Remarkably, the genes upregulated in DAPT versus vehicle treated *Neurog3*-null explants, revealed several acinar cell markers, such as *Ptf1a, Rbpjl, Cpa1* and *Cpa2*, which was also reflected in the heatmap of the corresponding genes mapped to the expression data from the E15.5 FACS sorted pancreas populations where the top genes are mostly expressed in the tip/acinar compartment (Figure 4C). However, comparing DAPT treated Neurog3-null versus wildtype, did not reveal tip/acinar genes to be upregulated (Figure 4C and Sup. Figure 5), and we therefore conclude that the propensity to upregulate acinar genes upon Notch inhibition is not restricted to the Neurog3-null population per se. A previous report also noted increased acinar differentiation after DAPT treatment of E12.5 wildtype explants [37]. Lineage tracing experiments are necessary to fully assess whether it is trunk progenitors or MPCs that respond to DAPT by upregulating acinar markers in our setup.

### Cbfa 2 t3, Ptf1 a, Gfi1, Gata 4 and Nkx 6-1 are Neurog3-Independent Targets of Hes 1

Next we compared genes bound by Hes1 in 266-6 cells with genes deregulated in the DAPT versus vehicle treated *Neurog3*-null explants and found 13 genes. Of these, only *Tm4sf1* was downregulated with DAPT treatment, while the remaining 12 Hes1 bound, derepressed genes include both of the master regulators of tip/trunk patterning *Nkx6-1* and *Ptf1a,* as well as other important transcriptional regulators such as *Gata4, Gfi1, Bhlha15* and *Cbfa2t3* (Figure 4D). Ptf1a, Gata4, Bhlha15, and Gfi1 are transcriptional regulators necessary for proper development of tip fate and/or full maturation of tip progenitors to enzyme producing acinar cells, processes that is suppressed by Notch signaling [15, 16, 38-45]. We show here Hes1 is binding these genes and that *Ptf1a, Gata4, Bhlha15*, and, *Gfi1* are all derepressed by DAPT treatment of Neurog-null explants, (5.26-fold, adj. p-value 0.036; 3.01-fold, adj. p-value 0.06; and 3.2-fold, adj. p-value 0.036 versus vehicle controls, respectively). Many of these genes are expressed in MPCs before becoming restricted to acinar cells during tip/trunk patterning and forced expression Ptf1a in MPCs is capable of inducing a tip cell fate, the same fate that is induced by loss of Hes1 [20]. This finding suggests that one function of Hes1 in MPCs and/or trunk cells is to prevent excessive tip cell allocation.

Intriguingly, our analysis also identified *Cbfa2t3 (*also known as Mgt16, Eto2 or Zmynd4) as a novel Hes1 target gene, bound in 266-6 ChIP-seq, and upregulated *ex vivo* in DAPT versus vehicle treated *Neurog3*-null explants (3.17-fold, adj. p-value 0.04). *Cbfa2t3* encodes a transcriptional corepressor interacting with e.g. N-ICD, E-proteins, and Gfi1 suggesting that it could be important for pancreatic cell fate decisions. Together, these data suggest that Notch signalling suppresses acinar development by Hes1-mediated repression of a set of transcription factors that act together to induce the acinar fate.

## Discussion

Here we have identified direct Hes1 target genes in pancreatic cells. Most notably, our ChIP-seq identification of genome-wide Hes1 binding sites revealed several transcriptional regulators involved in tip/trunk patterning. These genes were also derepressed in *Hes1*^−/-^ 266-6 cells and/or DAPT-treated embryonic pancreas explants, demonstrating that they represent *bona fide* direct targets of Hes1. By generating gene expression profiles of purified cell populations isolated from DBA-labelled E15.5 embryonic pancreas expressing lineage-specific fluorescent reporters we were able to map data from our subsequent experiments where Notch signalling and cell fates were perturbed. This provided a valuable framework for interpretation of the gene expression changes caused by Notch inhibition and Hes1 loss-of-function mutations.

Surprisingly, the presumed early (DBA^+^) and late (DBA^−^) endocrine precursors did not exhibit any significant differences in their transcriptomes but were, as expected, very different from the trunk and tip populations. They expressed the expected endocrine markers, and revealed a set of genes not previously reported to be enriched in the endocrine lineage. For the tip and trunk progenitors the differences were less pronounced, similar to observations by Benitez and co-workers [21], however, both trunk and tip progenitors showed enrichment of the expected markers. With the advent of single cell RNA-sequencing we anticipate that gene expression atlases like this will be a valuable additional resource due to its greater depth of transcript detection.

To gain insight into the direct molecular targets of the Notch pathway we perform Hes1 ChIP-seq, which provided 359 high confidence peaks overlapping between two biological replicates. We detected the classically defined C-site motif CACGCG [27] by Homer *de novo* motif search, which also extended the motif to ggCACGCGcc, which was found on the majority of peaks near promoters. Notably, we detected binding to the *Hes1* promoter, which in other systems enables an auto-repressive feedback loop necessary for the oscillatory expression of Hes1 [32, 46]. We found several Hes1 bound genes in the pancreas which were previously found in mouse embryonic stem cells by ChIP-on-chip analysis [32], including *Gadd45g, Neurog2, Ptf1a, Dll4* and *Ascl1*, revealing that Hes1 binds many of the same sites independent of cell type or tissue.

Surprisingly, we do not find *Neurog3* among the Hes1 bound genes in the acinar carcinoma derived 266-6 cell line despite that Hes1 has been shown to regulate and bind *Neurog3* in mouse pancreas and other tissues [6, 32, 34]. We speculate that the *Neurog3* locus would be epigenetically closed in tip/acinar cells, and this might be the reason for the lack of *Neurog3* binding, perhaps through DNA methylation of the CpG dinucleotide found in the Hes1 motif, which prevents binding [47]. In contrast, we would expect to find Hes1 binding to *Neurog3* in any cell that has potential to become endocrine Neurog3^+^ such as embryonic stem cells, MPCs, trunk progenitors, bile ducts [19], stomach and intestine [6, 48, 49]. Indeed, ChIP-seq for HES1 in human ES cells combined with mutational analysis reveals many genes in the endocrine lineage being bound and regulated by HES1 (see accompanying manuscript). Commitment to the endocrine lineage begins with a short, transient burst of expression of Neurog3 that brings on a cascade of transcription factors to regulate the differentiation to an endocrine hormone-producing cell that migrates from the duct epithelium to the islets of Langerhans. Several reports have shown that down-regulation of Notch signalling and consequently Hes1 allows de-repression of Neurog3 and commitment to the endocrine lineage [6, 8, 34]. However, to assess Neurog3-independent transcriptional changes upon Notch inhibition we sought to mask the secondary effects of endocrine differentiation by using *Neurog3*-null mouse explants. Despite high variability in the DMSO treated wildtype explants we were able to verify that DAPT treatment for 24h was sufficient to upregulate Neurog3 and the lack of endocrine marker expression in the *Neurog3*-null, confirmed the validity of our setup. We found that Notch inhibition have the potential upregulate tip/acinar markers both in wildtype and *Neurog3*-null explants, which is in line with previous observations [37, 50]. Qu and colleagues also find upregulation of acinar markers as well as insulin upon DAPT treatment of wildtype explants [37]. However, in that study DAPT was present throughout the culture period, which precludes direct comparison to our data. Tamoxifen-based tracing of “failed” endocrine precursors in *Neurog3*^CreERT/CreERT^ null-mutants, to duct and acinar cells before E12.5, but exclusively to duct cells after E12.5 [51, 52], suggests that cells activating the Neurog3-promoter lose the ability to become acinar cells after E13. However, when treating our Neurog3-null explants with DAPT we observe upregulation of numerous tip/acinar-expressed genes, suggesting that the *Neurog3* null cells at E12.5+3d explant culture do have the potential to become acinar. This is most likely explained by the delay in development that is consistently observed in explant systems [53].

Among the acinar markers genes upregulated in the Neurog3-null with DAPT treatment are *Ptf1a, Gata4, Bhlha15*, and *Gfi1*, which we also found to be bound by Hes1 in our ChIP-seq data. We show that Notch signalling controls multiple tip/acinar transcription factors necessary for tip fate allocation and maturation of acinar cells via Hes1 binding directly, not only to *Ptf1a*, but also the novel Hes1 target gene *Gata4*, both of which are upregulated with DAPT treatment independently of Neurog3. Hes1 binding to *Ptf1a* has been suggested before [18, 19], but has not previously been demonstrated by a ChIP-based assay. Ptf1a and Nkx6-1 are believed to be the main transcription factors driving tip/trunk patterning, acting in a mutually antagonistic fashion [13]. Notch signalling acts upstream of these transcription factors, but the exact mechanism remains incompletely understood. Constitutive Notch signalling in MPCs via *Pdx1* promoter-driven N1-ICD misexpression, favours an Nkx6-1^+^ trunk fate [13], while dnMaml1 mis-expression, as well as conditional *Mib1*-and *Hes1*-null mutations forces MPCs to adopt a tip fate [17, 20]. Notch has been proposed to activate *Nkx6-1* expression through direct binding of the NICD-Rbpj complex to a regulatory site in the *Nkx6-1* gene, which may then subsequently directly repress *Ptf1a* expression [13, 17]. Conversely, Ptf1a has been proposed to repress *Nkx6-1* indirectly. The data we present here adds to this picture by demonstrating that Hes1 can repress *Ptf1a* expression directly, possibly in cooperation with Nkx6-1 and by suggesting that Hes1 and Ptf1a may act together to repress *Nkx6-1* when MPCs initiate tip fate allocation. Nkx6-1 may then later be maintained in a repressed state in tip cells by other, perhaps epigenetic, factors. Moreover, the relatively low level of Nkx6-1 expression in the trunk progenitors, compared to β-cells [54, 55], may be explained by opposing inputs from NICD [17] and Hes1 (this study). Remarkably, our data suggest direct, Hes1-mediated repression of *Gata4, Bhlha15*, and *Gfi1*. *Gata4* and *Gata6* are both expressed in MPCs and simultaneous loss of these genes in mice results in pancreas agenesis while loss of Gata4 alone causes deficiencies in acinar development [44, 56]. Although, Gfi1 is best known for its requirement for intestinal goblet cell differentiation, Gfi1 is also expressed in embryonic pancreas from E11.5 and becomes restricted to tip/acinar cells later in development and knockout of Gfi1 results in exocrine dysplasia through defects in centroacinar cells [42]. The direct repression of *Ptf1a, Gata4, Bhlha15*, and *Gfi1* by Hes1 suggests that Notch signalling suppresses acinar differentiation and maturation through multiple repressive “handles”. The extensive overlap between Hes1 peaks and published Ptf1a ChIP-seq peaks on distal sites (>2 kb away from TSS) is remarkable in this context. By immunoprecipitation we showed that endogenous Ptf1a protein can pull down Hes1, extending previous observations with overexpressed, epitope-tagged proteins [57]. The trimeric Ptf1 complex includes Ptf1a, and E-protein, and Rbpj [58, 59], which intriguingly also binds to the *Hes1* promoter [36, 40, 60]. Our data showing Hes1 and Ptf1a binding to neighbouring sites at a large number of loci makes it tempting to speculate that they can be bound simultaneously to form a repressive complex providing a means to convert the transcriptional activator Ptf1 into a repressor as well as possible feedback loops to the Notch pathway.

Notably, we find the transcriptional repressor Cbfa2t3 (also known as Mtg16, Eto2 or Zmynd4) to be a Hes1 bound gene, which is upregulated upon Notch inhibition in the *Neurog3*-null explants. Based on the literature Cbfa2t3 appears to be involved in numerous aspects of Notch mediated cell fate choice and bHLH-induced lineage commitment. The Cbfa2t3^−/-^ mouse has defective crypt proliferation and goblet cell differentiation, which may also implicate another Hes1 target gene, *Gfi1*, identified here. However, its role in the pancreas has only been studied in *Xenopus* where it is downstream of Neurog3 and is necessary for differentiation to insulin producing cells, ostensibly by downregulating Neurog3 [41, 61-64]. In neural progenitors Cbfa2t3 it interacts with Ascl1 and Neurog2 and forms a negative feedback loop that represses Ascl1 and Neurog2 and it might act similarly in the pancreas on Neurog3 [65]. Further investigations in the role of Cbfa2t3 are necessary to shed light on its putative role in pancreatic cell fate choices.

Taken together, we performed the first ChIP-seq on endogenous Hes1, observed substantial overlap with Ptf1a on distal elements and we find Notch signalling control acinar differentiation including acinar genes Ptf1a, Gata4, Gfi1 and trunk/endocrine gene Nkx6-1 to be bound and repressed by Hes1 in a Neurog3 independent manner. This shows the specific means by which Notch signalling is controlling key transcription factors and thereby cell fate choice.

## Materials and Methods

### Mouse lines

Hes1-EGFP, Neurog3-RFP, Neurog3^tTA^, Ptf1a^YFP/+^ mouse lines were described previously [66-69]. Animals were maintained in adherence to guidelines issued by the European Convention for the Protection of Vertebrate Animals used for Experimental and other Scientific Purposes (ETS 123).

### E15. 5 FACS sorted populations

Mice were genotyped for heterozygosity by qPCR compared to wildtype and known homozygous by test-crosses. qPCR was performed according to manufacturer’s instructions using ROCHE UPL probes as below on a Lightcycler 480II and compared to control locus: Ubc Promoter.

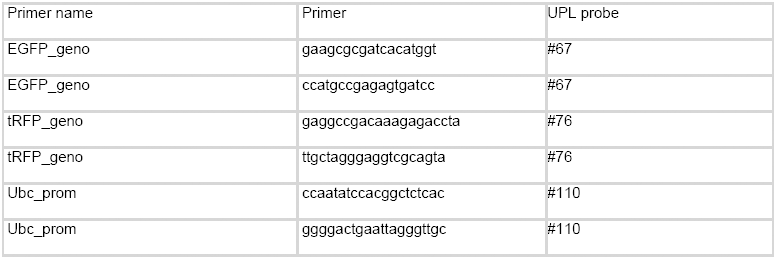

Hes1-EGFP homozygous were set up with homozygous Neurog3-RFP females for timed matings and harvested at E15.5 for fine dissection of a pool of ventral and dorsal buds from 5 embryos to MEM (Gibco) on ice. To each tube added DNAse I (Invitrogen, 1:300) and 1:400 Liberase TL (of 117U/mL) and incubated 20 minutes at 37°C on a thermoshaker, 950rpm, with trituration every 3-4 minutes. Spin down 500g 3 minutes at 4 °C. Then resuspend in PBS with 1:20 Trypsin-EDTA 0.125% (Gibco) and DNase I as above. Incubate 10 minutes 37 °C shaking and trituration as above. Spin as above and wash pellet in 10% FBS in PBS. Spin as above and added DBA-biotin (Vector-labs) 1:200 in 10%FBS in PBS, rotating at 4°C for 10 minutes. Washed in 10%FBS in PBS, added Straptavidin-APC-Alexa750 1:1000 rotating 15 minutes at 4 °C. Washed in 10% FBS in PBS, spin as above and resuspended in 10%FBS in PBS + DAPI.

Fluorescence activated cell sorting was done on BD Bioscience ARIA II. For Hes1GFP mated with Neurog3-RFP, gating GFP+, then sorting GFP+DBA+, GFP+DBA+RFP+ and GFP+RFP+ as indicated in Figure 1. Sorting was done directly to RLT+ for RNA purification with MicroElute RNeasy Plus (Qiagen). Ptf1aYFP/+ embryos were dissected and dissociated as above and sorted directly to RLT plus and purified with MicroElute RNeasy Plus (Qiagen).

### Neurog 3 t TA explants E 12.5+4d

Heterozygous crosses of Neurog3^tTA/+^ were set up for timed matings and at E12.5 dorsal pancreata were dissected and adhered to fibronectin coated slides (BD Bioscience) in DMEM (Gibco) with 20% FBS. Changed media once daily and 24h before harvest added 10μM DAPT (Sigma, D5942) or DMSO as vehicle control. Explants were harvested in RLT and purified with MicroElute RNeasy Plus (Qiagen).

### Microarray of E 15.5 FACS populations and Explants

Quality of the RNA was checked by Bioanalyser pico kit (Agilent) and using 35 ng total RNA was subjected to amplification and hybridisation was done using Low Input Quick Input Amp Kit (Agilent) and loaded onto Mouse GE 8x 60k microarrays (Agilent).

### Hes 1 Ch IP in 266 −6 cell line

266-6 cells were grown in DMEM (Gibco) supplemented with 10%FBS on plates coated with 0.01% Matrigel (growth factor reduced) in PBS, incubating for 1 h at room temperature.

### Ptf1 a immunoprecipitation

For immunoprecipitation, cells were seeded at 1.2 mio cells in a 10cm plate, 3days later harvested in PBS on ice, scraped and spun down 500g 4 °C then resuspended pellet in RIPA buffer and lysed 20 minutes on ice. Then sonicated 7 ×; 30sec ON/OFF on Bioruptor (Diagenode), spun full speed 30 minutes 4 °C. For input 2% was set aside. Antibodies used for IP: 2μL Rb IgG (Cells Signaling), 1μL Hes1 Rb H140-X (Santa Cruz), 1 μL Rb Ptf1a (BCBC-2432A). Rotated over night at 4 °C, then added 30μL dynabeads (Thermo) and incubated 2.5h further. Washed 3x in IP buffer and eluted in 2x sample buffer. Western blot as below.

### CRISPR-Cas 9 mediated Hes 1 ^−/-^ cell line

Crispr gRNA design was taken from the Gecko Library (Hes1,MGLibB_24024) cloned into px330 according to protocols by Feng Zhang lab and co-transfected with an IRES-GFP plasmid into 266-6 cells with lipofectamine 200 according to manufacturer’s instructions. 48h later sorted GFP+ cells and seeded at clonal density. Picked clones and expanded.

### Genotyping 266-6

PCR genotyping with Phusion polymerase (Thermo) using Tm 60 °C.

Forward primer: AAGTTTCACACGAGCCGTTC

Reverse primer: CATTTCACCCCGAGGTTTTA

PCR Product was then sanger sequenced to verify indel formation and subsequently knock-out was verified by western blot.

### Western Blot

266-6 clones (#1=wildtype, #4 and #5 Hes1 knock-out), were treated with DAPT overnight or vehicle control DMSO. 2h washout samples were first treated with DAPT overnight then media was replaced by normal growth medium without DAPT. Antibodies used for western blot: α-Nkx6-1 (BCBC 2022, 1:1000, Mouse), α-Tubulin (, 1:5000, Rat), α-N1-ICD (Cell Signaling, 1:1000, Rb), α-Hes1 (Santa Cruz H140-X, 1:2000, Rabbit). Secondary antibodies were from Jackson laboratories, for tubulin we used α-Rat Cy5 and for the rest HRP conjugated corresponding to primary antibody species.

### RT-qPCR

266-6 clones were harvested in RLT plus and purified with Mini RNeasy Kit (Qiagen). Quality and quantity were measured on a Nanodrop 2000 and 1μg of total RNA was used for RT reaction with Superscript III (ThermoFisher) according to Manufacturer’s instructions using a mix of Random Hexamers and Oligo-dT primer in a 20 μL reaction. For qPCR the UPL system from Roche was used according to manufacturer’s instructions in 10μL reaction volume in 384 well format with following primers:

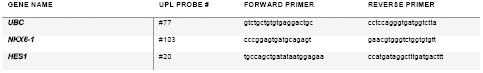

### Chromatin Immunoprecipitation (Ch IP) and libraries for sequencing

Since the experiment was setup for both Notch1 and Hes1 ChIP, 266-6 cells were transfected with N1-ΔECD, a γ-secretase dependent but ligand independent Notch1 previously published [70] which has a 6x-myc tags. Cells were transfected in 10cm plates using using 30μL Lipofectamine 2000 in 600μL optiMEM + 20μg plasmid DNA according to manufacturer’s instructions, changed media the next day to either DAPT containing or vehicle control DMSO. Cells were fixed in DSG for 30 minutes for protein-protein interactions, then fixed 1% Formaldhyde in PBS (freshly made from Amersham MeOH-free 16% Formaldehyde) for 10 minutes. Stopped fixation in 0.125M Glycine for 5 minutes and washed twice in PBS. Then did nuclear enrichment in 0.5% SDS ChIP buffer for 5 minutes and discarded supernatant after spinning. Lysed in RIPA buffer with protease inhibitors for 10 minutes on ice. Sonicated 7 ×; 30seconds ON/OFF, then spun full speed for 20 minutes. Used 40μg chromatin for each ChIP with 4μg antibody was used: αIgG (Cells Signaling), rabbit, αHes1 Rb H140-X (Santa Cruz), αMyc (A14). ChIP samples tumbled 3.5h at 4 °C. Added Protein G Dynabeads and tumbled for 45 minutes, washed in RIPA + inhibitors twice, then once in TE buffer moving to a fresh eppendorff tube and eluted in Elution buffer containing 1%NaHCO_3_, 1% SDS and proteinase K in PBS for 2h at 56°C and then overnight at 65°C. Cleaned up ChIP’ed DNA with ChIP Clean and Concentrator kit (Zymo) and quantified on a Qubit (ThermoFisher) for sequencing. Multiplex libraries were built using a published protocol developed in the Ido Amit lab [71] and sequenced on an Illumina HiSeq.

### Ch IP-sequencing bioinformatics analysis

Mapped reads with Bowtie2 to mm10 and tracks for the IGV browser were made using DeepTools. Peakfinding was performed using MACS2, subtracted Encode Blacklist and did de novo motif finding was done using Homer using 200 base pairs surrounding the summit of the peaks. Ptf1a, Rbpj, Rbpjl, and Foxa2 ChIP-seq was previously published by [14, 36, 40], was mapped to mm10 as above and heatmap of ChIP-seq signal on Hes1 peaks was made using Deeptools Computematrix function, defining the promoter region as within 2kb of TSS. GREAT software (http://great.stanford.edu/public/html/) was used for database enrichments showed in Figure 2E. For all other annotations ChIP-seeeker was used [72]. IGV software was used for visualisation of ChIP-sequencing tracks.

### Microarray gene expression data analysis

Limma package was used for both microarray experiments as detailed in userguide for quality check, hierarchical clustering, MultiDimensional Scaling plot and to find differential expression. Visualised within R environment with packages ggplot, ggrepel, pheatmap, and Superheat package for heatmaps with kmeans clustering using WardD2 method when applicaple.

Gene set enrichment Analysis was done using GSEA Java application loading the microarray datasets and finding enrichment comparing to C2 database of MsigDB. For Neurog3tTA explant microarrays the data from E15.5 FACS population studies described above providing specific genes expressed in Acinar, Endocrine and trunk/duct progenitor cell types were uploaded as custom gene sets and compared alongside C2 database.

